# Cohesion in male singing behavior predicts group reproductive output in a social songbird

**DOI:** 10.1101/2021.11.20.469403

**Authors:** Ammon Perkes, H. Luke Anderson, Julie Gros-Louis, Marc Schmidt, David White

## Abstract

All social groups require organization to function optimally. Group organization is often shaped by social ‘rules’, which function to manage conflict, discourage cheating, or promote cooperation^1–5^. If social rules promote effective social living, then the ability to learn and follow these rules may be expected to influence individual and group-level fitness. However, such links can rarely be tested, due to the complexity of the factors mediating social systems and the difficulty of gathering data across multiple groups. Songbirds offer an opportunity to investigate the link between social rules and reproductive output because most of their social interactions are mediated by song, a well-studied and readily quantifiable behavior^6,7^. Using observations from 19 groups of brown-headed cowbirds (*Molothrus ater*) studied across 15 years, we find evidence for a previously undocumented social rule: cohesive group transitions between dominance- and courtship-related singing. Comparing across groups, the degree of cohesion in male singing behavior predicts the reproductive output of their group. Experimental manipulation of group structure via the introduction of juvenile males to captive flocks reduced group cohesion and adult male reproductive success. Taken together, these results demonstrate that cohesion in group behavioral states can affect both individual and group-level reproductive success, suggesting that selection can act not only on individual-level traits, but also on an individual’s ability and opportunity to participate effectively in organized social interactions. Social cohesion could therefore be an unappreciated force affecting social evolution in many diverse systems.

## Results and Discussion

In diverse animal taxa, there exists clear structure in social interactions, which are reliable, consistent, and often ritualized. While in some systems these rules of interaction may be imposed from the top down by leaders, often they emerge as a product of individuals interacting in complex systems. Testing the group-level effects of these rules requires comparing fitness across multiple groups. Here, using a 15-year dataset in brown-headed cowbirds (*Molothrus ater*), we show that group-level cohesion in song use is linked to variation in reproductive output across groups.

### Male cowbirds are cohesive in their use of singing strategies

In brown-headed cowbirds, males use song for two primary purposes: (1) to establish male dominance hierarchies via male-directed countersinging; and (2) to attract and maintain pair bonds via female-directed courtship singing^8^. In captive cowbirds housed in large, outdoor aviaries, we observed that groups of males collectively transitioned, over the timescale of minutes, between periods of primarily male-directed song and periods of mostly female-directed song (Figure 1a-b; supplemental video 1). Unlike the tightly synchronized and/or cooperative displays observed in some organisms^9–11^, male cowbirds court females independently but tend to sing to their respective pair bonds at similar times, generating a group-level behavioral organization of courtship.

**Figure 1.**
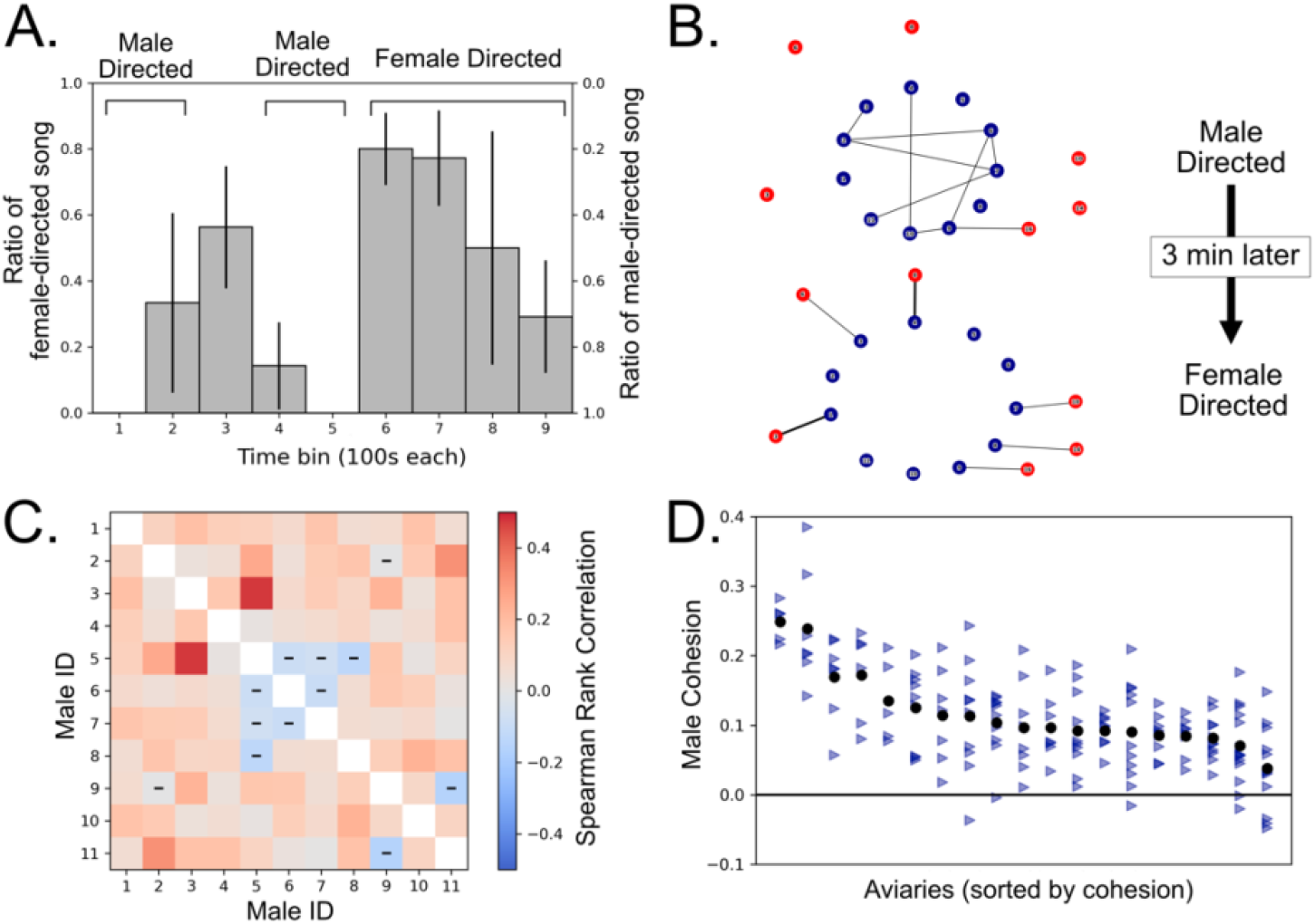
Social groups are cohesive across time. **A. Long-term transition between song strategies.** In this plot, the heights of the bars represent the proportion of female directed song, or equivalently 1 minus the proportion of male-directed song, over the course of 15 minutes. Over the plotted time course, males transition between male- and female-directed song twice, with a periodicity of roughly five minutes. **B**. **Network representation of singing behavior.** These network plots show one transition from male-directed to female directed song at the level of individual behavior. Males (blue nodes) sing to other males and females (red nodes) during two 100s time bins occurring 5 minutes apart. Edges between nodes represent at least one song, with the thickness of lines depicting the number of songs sung. Females who were not sung to in either bin were omitted for clarity. **C. Pairwise cohesion for a single group.** This heatmap depicts the pairwise cohesion between all males within a single group. Negative correlations (blue) are marked with hashes. **D. Aviary-level variation in cohesion.** This plot depicts the variation in cohesion across aviaries, showing that the majority of aviaries show significant levels of cohesion. Each black dot shows the mean aviary cohesion for a single group, produced by averaging all pairwise correlations (the values depicted in C). The columns of blue triangles represent the mean cohesion for individual males (i.e., the row averages from C) from the same group. Columns (i.e., groups) are sorted by mean aviary cohesion. All but one aviary (the rightmost one) was significantly more cohesive than random, confirming that the cohesive patterns observed in A and B are statistically significant.

We termed the coordination of these group-level transitions between behavioral states “cohesion,” which we quantified as the degree to which singing behavior was temporally correlated among males in a group. Specifically, we divided the breeding season into short time bins and calculated the percentage of songs directed to females within each bin. This yields a group distribution of the proportion of songs directed to females. While independent male behavior would result in an approximately binomial distribution, cohesive singing behavior should increase the number of bins with high male- or female- directed song (Figure S1a). We compared the observed distribution with a random distribution (generated by shuffling the observed bins for each male) to determine whether real groups exhibited more cohesion than expected by chance. In 18 of 19 aviaries analyzed, distributions were significantly skewed towards periods of high male-directed song and periods of high female-directed song, compared to their shuffled distributions (Figure S1b, KS test, p<0.05, Table 1). Thus, at the group level, males are more cohesive in their singing behavior than can be explained by chance interactions.

To better compare cohesion across groups, we devised a standardized metric to calculate the overall level of cohesion for each group by correlating each male’s singing behavior (i.e., the proportion of songs sung to females in a given time bins) with that of each other male in that group (Figure 1d, see Methods). These values were then averaged to calculate the overall level of cohesion among males in each group, while also providing an estimate of each male’s mean individual cohesion (Figure 1d). To evaluate the statistical strength of cohesion, we again generated a random distribution of cohesion by shuffling the order of each male’s binned behavior and repeating the calculation of mean pair-wise cohesion (see Methods). Here, 18 of 19 aviaries were significantly more cohesive than their respective shuffled distributions (Figure 1d). Taken together, these results demonstrate that males behave cohesively in their use of female- and male-directed songs, suggesting group coordination of singing.

### Male cohesion predicts group-level reproductive output

While all aviaries displayed cohesion in group singing behavior, we observed significant variation in the degree of cohesion between aviaries (one-way ANOVA, 6.55, p<0.0001). We then tested whether this variation in cohesion correlated with reproductive success. Female cowbirds lay readily in captivity when provided with mock host nests and exhibit considerable variability in egg output^12^, providing an opportunity to quantify differences in reproductive fitness among aviaries. We calculated the reproductive output score of each aviary (normalized to account for variation in sampling period and number of females, see Methods). We found that more cohesive aviaries had significantly higher reproductive scores (Figure 2, r = 0.43, CI: 0.01-0.83, p=0.046), revealing a link between the strength of cohesion and the reproductive fitness of the group.

**Figure 2.**
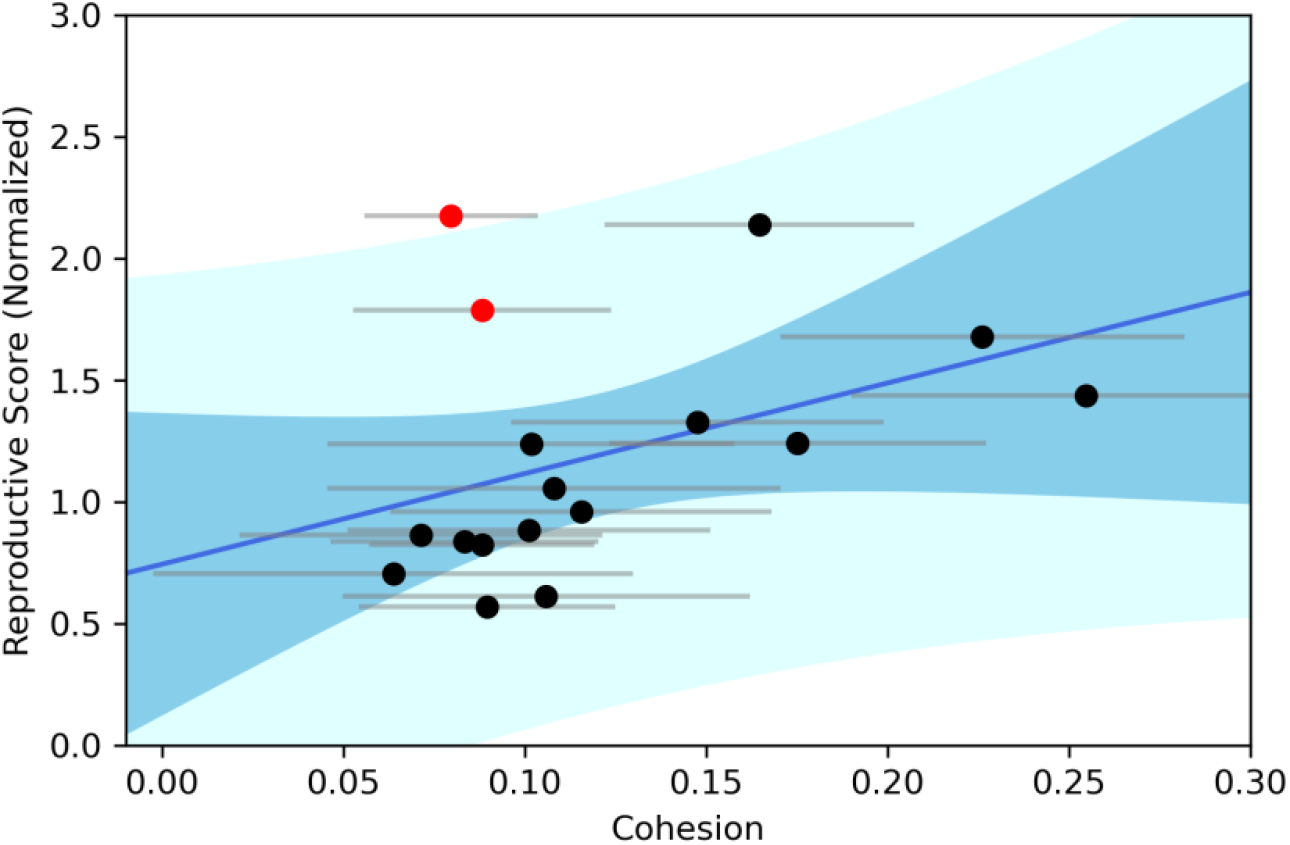
Aviary cohesion predicts egg production. The scatter plot shows the normalized egg production of each aviary as a function of aviary cohesion. The blue line shows the estimated effect based on linear regression. Aviary cohesion significantly predicted total egg production (r=0.42, p=0.046), showing that more cohesive groups produce greater numbers of eggs. Cohesion is calculated as the mean across all pairwise interactions (equivalent to Figure 1d), and egg production is normalized based on the number of females present and the date ranges observed in order to produce the reproductive score plotted on the y-axis. Horizontal error bars represent the variation among males within a given aviary. The light blue and dark blue shading represent the 95% confidence interval of the regression and the effect strength, respectively. Red dots denote the two aviaries in which pair bonds were maintained from the prior year. These dots did not fit the overall trend, suggesting that prior pair bond experience may have masked any effects of cohesion in these two groups.

This effect of cohesion was strong: the most cohesive group produced 44% more eggs than average and the least cohesive group produced 30% fewer than average. Despite the strong overall trend, two aviaries from 2009 stood out as exceptions (Figure 2, red dots). On further investigation, these were the only two aviaries containing repeated male-female pair bonds from the previous year. Greater pair bond familiarity may have increased egg production^13^, separate from the effect of cohesion, or it may be that the effect of cohesion is particularly strong during pair bond formation, with established pair bonds being more robust to group effects.

### Male cohesion increases reproductive output by increasing the number of females participating in reproduction

Our prior result demonstrates a link between cohesion and reproductive output, but does not reveal how cohesion might influence the level of reproduction within the group. Increased reproductive output in more cohesive groups could be driven by greater fecundity per egg-laying female, by an increase in the number of egg-laying females, or both. To distinguish between these possibilities, we tested whether each of these two measures was correlated with reproductive output across groups. We found a strong correlation between aviary-level egg production and the proportion of females participating in courtship (r=0.55,p=0.023) but no correlation between overall egg production and number of eggs produced per participating female (r=−0.02), suggesting that variation in reproduction was primarily a function of the number of egg-laying females. With regards to cohesion, we similarly found no effect of cohesion on the average number of eggs per participating female (r=−0.22, p=0.46), but cohesion did appear to predict the proportion of females who laid at least one egg (r = 0.55,p=0.023). These findings indicate that cohesion-associated change in aviary productivity was driven primarily by an increase in the number of females participating in reproduction rather than an increase in productivity by participating females.

### Individual-level cohesion predicts pair bond formation but not fecundity

While cohesion predicted egg production at the group level, we wanted to test whether a male’s relative cohesion within a group affects individual success. Cowbirds form stable pair bonds which persist across seasons, with each female receiving the majority of songs from a single male^14^. While each female tends to have one distinct pair bond male, individual males can have multiple pair bonds with different females, thus a male’s reproductive output depends both on the number of pair bonds acquired and the number of eggs produced by each pair bond. We quantified the reproductive output of individual females by video-tracking their visits to artificial host nests in 13 of the aviaries. Prior work has shown that extra-pair paternity is very rare in aviaries^14^, thus we assigned all of a female’s eggs to her respective pair bond to infer male reproductive fitness.

Using a network approach (see Methods), we found that within aviaries, greater male cohesion (with respect to other males), predicted greater total reproductive output of his associated pair bond(s) (coef = 0.850, CI: 0.247-1.453, p=0.006). The strength of individual cohesion on egg production suggested that, in general, cohesion was a strong predictor of bonding with a productive female and a fruitful reproductive strategy for males at the individual level.

### Alternate Strategies

Despite the individual and group-level benefit of cohesion, we found evidence for alternative strategies among some males. Within these aviaries, male reproductive success was a function of the eggs produced by the primary pair bond (effect = 0.85, CI: 0.75-0.94), as well as the total number pair bonds acquired (effect = 2.23, CI: 1.83-2.65). Obviously, males with pair bonds obtained more eggs than those without, and individual cohesion was a strong predictor of obtaining a pair bond (see Figure 3 above), but within successful males—those who managed to obtain at least one egg laying female as a pair bond—higher cohesion was a poor predictor of increased reproductive output (effect =−0.007, CI: −0.928-0.914), and in 8 of 13 aviaries, a greater number of eggs actually appeared to be associated with decreased cohesion (mean r = - 0.34 +− 0.19). This suggests that, while cohesion is important for groups as a whole and for individuals forming at least one successful pair bond, some males obtain higher reproductive output while behaving less cohesively. This apparent incentive for decreased cohesion among some successful males may explain why variation in cohesion persists across males and groups.

**Figure 3.**
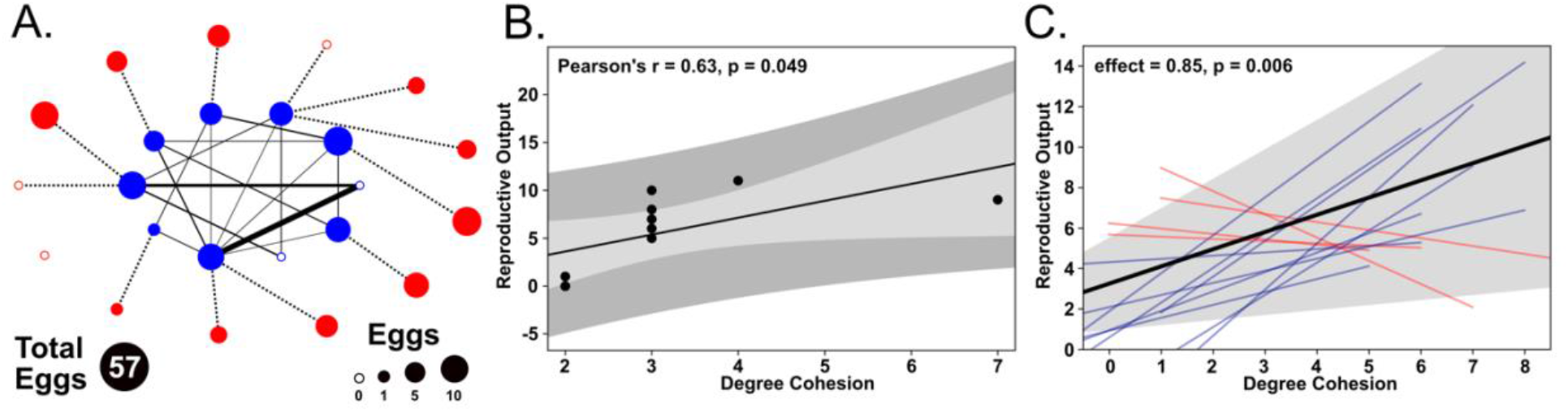
Individual cohesion predicts egg production. **A. Cohesion network and egg production.** This network plot depicts the strength of cohesion and eggs acquired by each individual male (blue interior nodes) and female (red exterior nodes). The male:male edges in this network represent the strength of pairwise cohesion, as in Figure 1C, with thicker lines representing a higher correlation. The dotted lines between male and female nodes denote pair bonds. The size of nodes (for both males and females) is proportional to the number of eggs produced, with the smallest nodes yielding 0 eggs (also denoted by open circles). Correlations below 0.1 were omitted for clarity. **B. Reproductive output as a function of cohesion from a single aviary.** This plot shows a regression for the same group of birds in A, between the reproductive output (denoted by size in A) and individual cohesion. Here individual cohesion is measured as degree, i.e. the number of male:male edges for each male from the network in A. Males with a large number of cohesive links (3 or more) sired higher numbers of eggs within this aviary. **C. Reproductive output as a function of cohesion for all aviaries**. The black line shows the effect of individual cohesion across all aviaries (coef = 0.85,p=0.006), based on a linear mixed model with aviary as a random intercept. The gray shading showing the 95% confidence interval as predicted by the model. The colored lines show simple linear regression across each aviary, with blue lines showing a positive correlation and red lines showing a negative one. These results suggest that individual cohesion is associated with higher individual reproduction.

### Experimental disruption of cohesion reduces egg production: a case study

Finally, we attempted to determine whether it was possible to experimentally decrease cohesion and influence reproductive success. To do so, we reexamined data from a prior experiment in which we manipulated the age-class composition of two cowbird flocks over two years by either adding or removing juvenile males. This disruption to the social environment by the addition of juveniles caused a dramatic decrease in egg production^15^. However, the mechanism of the effect has remained a 15-year mystery. Our original analyses failed to find any change in the adult male behavior that could account for the reproductive decline in the group’s females. We were left merely to speculate that “the social chemistry that results from interactions within a group affects reproductive stimulation^15^” (Gros-Louis et al, 2006, p. 232).

Based on findings in our current study, we suspected that changes in cohesion might explain the effect observed in this experiment. Previous work has shown that the singing patterns of juvenile males are highly variable and unstructured compared to adults^16^, which could negatively impact group-level cohesion. In the first experimental flock, after one year of normal (adult male) conditions, in the second year we had replaced a subset of adult males with the same number of juveniles (Figure 4a). The addition of juvenile males indeed disrupted the groups: reproductive success of the resident adult males was lower the year juveniles were present. This effect was attributable to a dramatic decrease in female egg output in the groups where juvenile males were present (71 compared to 28 eggs). Applying our cohesion analysis to these males’ singing patterns, we observed that the addition of juvenile males was in fact associated with a drop in cohesion (Figure 4a), both for the group as a whole (decreasing from 0.12 to 0.07), and for individual males that remained in the group (decreasing from 0.14 in year 1 to 0.07 in year 2). Even more strikingly, we observed the emergence of ‘discordant’ behavior, i.e., negative correlations between males (visualized in Fig 4 as red lines) in 11% of all interactions (compared with just 1% in the adult aviary). These discordant interactions reflect the opposite of pairwise cohesion—one male consistently singing to females while the other is singing to males. This same pattern of discord was observed in the second flock, where we performed the opposite manipulation: juvenile males were added for the first year but were replaced with adult males in the second year (Figure 4b). Here again cohesion was low (.06), and discord was relatively common (8% of interactions in the juvenile aviary compared to 4% with adults), and reproductive output was again lower with juveniles present (25 eggs vs 59 eggs). Interestingly, in this second aviary, cohesion failed to recover when juveniles were replaced with adults (0.05 in the second year vs. 0.12 in group 1 adults), suggesting that cohesion and discord may act independently to influence reproductive output, or perhaps are simply two different facets of the same underlying organizational pattern. In aviaries with juveniles present, discord was specific to interactions involving juveniles (100% in group 1 and 83% in group 2).

**Figure 4.**
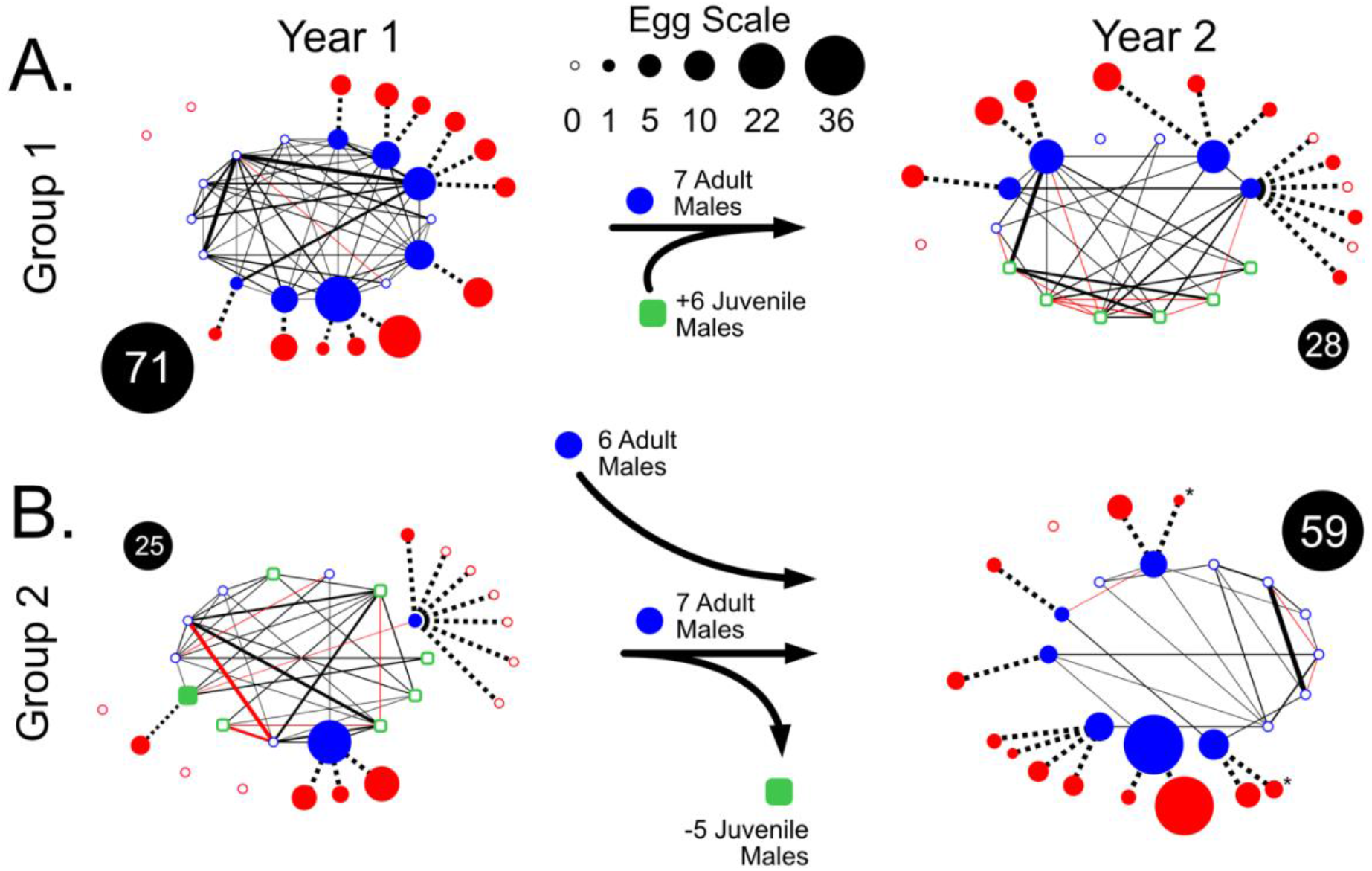
Experimentally disrupting cohesion reduces egg production. As in Figure 3, these plots depict the cohesion networks from two experimental aviaries across two years. In **A, juveniles were added in the second year**, replacing 7 adult males (blue nodes), while in **B, juveniles were removed in the second year**, and replaced with adults (who were brought from group 1). The presence of juveniles (green squares) is accompanied by lower egg production from the aviary (depicted in black circles), as well as lower adult male reproduction (shown as the area of blue nodes, see egg scale above). Juveniles were also accompanied by low average cohesion, as well as the large number of discordant interactions (Spearman’s rank coefficient < −0.1, marked in red). Taken together, this suggests that the presence of juveniles reduces reproductive output for males present, due to their disruption of normally cohesive behavior. As in Figure 3, the lines between male nodes show the Spearman’s rank coefficient of male singing behavior, with the thickness depicting the magnitude of the correlation. Dotted lines to females (red nodes) depict pair bonds, based on singing behavior and microsatellite analysis of the eggs to assign parentage. One female in Group 2, year 2 (marked with *) produced 1 egg from her non-pairbond male, thus she is plotted twice. Male positions are maintained across aviaries (i.e. the 7 adult males in Group 1, year 2—as well as the lower 6 males in Group 2, year 2—are plotted in the same positions as in Group 1, year 1).

The re-examination of this experiment provided several important insights. Critically, it suggests that group organization can be experimentally manipulated using network composition. In addition, it shows that cohesion is not a function of social contagion or social facilitation, as juveniles appear to lack the social skill necessary to generate cohesive networks, resulting in the disruption in the effectiveness of group organization and loss of reproductive output. Cohesion may therefore be the ‘social chemistry’ we posited 15 years ago.

## Conclusions

Our results reveal a novel form of coordination in a social species—cohesive transitioning between periods of competition and courtship—that provides a reproductive benefit to the group. There are many possible mechanisms by which cohesion could influence reproductive output. For example, cohesion may strengthen the efficacy of male signals, as has been demonstrated in other systems^17–19^. However, this seems unlikely in cowbirds, given the variation in song structure across males^20^ and the lack of precise display synchrony in time or space. Alternatively, cohesive behavioral transitions may increase reproductive success by generating a patterned social environment, providing predictable opportunities for females to eavesdrop on intrasexual competition and evaluate potential mates^21–23^ while also allowing for periods of stable courtship with reduced interference from other courting males^24–27^. The latter effect may be particularly important, as an ordered, reliable sequence of courtship behaviors can be vital for allowing males and females to progress towards effective reproduction^28^. We also note that, although we explored cohesion in male courtship behavior, female behavior may be equally important in shaping the organization of the group^29–31^.

Regardless of the specific mechanisms involved, a fitness benefit for cohesion *per se* has important implications for understanding how selection can act on a variety of other social species. If building and maintaining cohesion among group members is a skillful, long-term endeavor, as suggested by the behavior of juveniles in this study, then there may be high costs to leaving a cohesive group to become established in a new one. Cohesion may also shed light on mating system evolution. For example, pair bonding and social monogamy, in addition to providing direct benefits at the individual level, may facilitate group-level cohesion by reducing intrasexual competition once pair bonds are established. Pair bonds provide independent channels for courtship, allowing many pairs to court simultaneously without interference, however, the benefits of cohesion could also be realized in other mating systems: males in lek-mating species aggregate in specific locations to display to females^32^ , creating the opportunity for cohesive courtship. Numerous hypotheses have been proposed to explain the evolution of lekking^33–35^, but the possibility that behavioral cohesion at lek sites may yield group-level fecundity benefits has received little attention. While the above examples discuss direct benefits of cohesion for reproduction, the group-level effect of cohesion may be even stronger in systems^36^, including humans^37,38^ and other primates^39^, where cohesion provides indirect benefits to fitness by increasing a group’s ability to perform complex tasks.

While group-level selection remains controversial^40–42^, cohesion is a uniquely group-level phenomenon that correlates positively with reproductive output, thereby providing a potential mechanism by which higher-order selection could influence the evolution of social behavior. Importantly, the ability to manipulate cohesion provides an opportunity to experimentally test how selection operates at multiple levels. This work demonstrates the vital need for future research, both theoretical and empirical, to understand the ways in which group cohesion may impact individual- and group-level fitness outcomes in a wide range of social species.

## Methods

### Subjects and Housing

Male and female cowbirds were wild-caught in Montgomery County Pennsylvania (2002-2010), and in Ontario, Canada (2017) and housed in large (18.3 × 6.1 × 3.7 m) aviaries. Birds were maintained in similar conditions for all experiments (although specific flock distribution and bird densities vary, see Table 1). Aviaries contained artificial nests in which we placed faux white host “eggs” to stimulate laying. Captured individuals were housed in aviaries for at least 6 months prior to behavioral observations to allow time to habituate to captivity. Birds were fed a modified version of the Bronx Zoo Diet for omnivorous birds. Both food and water were provided ad libitum.

**Table 3.1.** Overview of aviaries. Specific values for each aviary. KS P-values below 0.0001 are expressed as 0.o001. Asterisks (*) denote aviaries for whom we could not assign parentage. Crosses (†) denote aviaries that ended before the laying season (these are only used for measuring cohesion)

| Aviary Name | Start Date | End Date | Males | Females | EggScore | Cohesion | KS Score | KS Pvalue |
| --- | --- | --- | --- | --- | --- | --- | --- | --- |
| PH1-2006* | 03-15-06 | 05-11-06 | 5 | 12 | 1.678 | 0.226 | 0.131 | **0.0001** |
| PH2-2006* | 03-22-06 | 05-11-06 | 14 | 13 | 0.705 | 0.064 | 0.055 | **0.037** |
| PH3-2006* | 03-15-06 | 05-11-06 | 5 | 11 | 2.140 | 0.165 | 0.084 | **0.0003** |
| PH4-2006* | 03-15-06 | 05-11-06 | 16 | 13 | 0.863 | 0.071 | 0.053 | **0.034** |
| PH1-2007† | 03-09-07 | 04-24-07 | 10 | 27 | Nan | 0.091 | 0.059 | **0.014** |
| PH2-2007† | 03-09-07 | 06-06-07 | 8 | 15 | 0.569 | 0.090 | 0.051 | **0.0006** |
| PH3-2007 | 03-09-07 | 06-05-07 | 8 | 19 | 0.824 | 0.088 | 0.044 | **0.002** |
| PH4-2007 | 03-09-07 | 04-24-07 | 11 | 24 | Nan | 0.104 | 0.038 | 0.305 |
| PH1-2008 | 03-26-08 | 06-06-08 | 10 | 14 | 0.883 | 0.101 | 0.051 | **0.002** |
| PH2-2008 | 05-10-08 | 06-06-08 | 7 | 9 | 1.327 | 0.147 | 0.076 | **0.005** |
| PH3-2008 | 05-10-08 | 06-06-08 | 9 | 10 | 0.959 | 0.115 | 0.052 | **0.037** |
| PH4-2008 | 03-26-08 | 06-06-08 | 11 | 17 | 0.838 | 0.083 | 0.049 | **0.002** |
| PH6-2009 | 04-22-09 | 06-02-09 | 10 | 13 | 2.176 | 0.080 | 0.044 | **0.004** |
| PH7-2009 | 05-28-09 | 06-09-09 | 8 | 12 | 1.239 | 0.102 | 0.063 | **0.002** |
| PH8-2009 | 04-22-09 | 06-02-09 | 10 | 13 | 1.788 | 0.088 | 0.054 | **0.0003** |
| PH6-2010 | 05-03-10 | 06-11-10 | 8 | 11 | 1.242 | 0.175 | 0.115 | **0.0001** |
| PH7-2010 | 04-10-24 | 06-11-10 | 9 | 10 | 1.436 | 0.255 | 0.098 | **0.0001** |
| ON1-2017 | 05-22-17 | 06-28-17 | 11 | 11 | 0.613 | 0.106 | 0.098 | **0.0001** |
| ON2-2017 | 05-21-17 | 07-04-17 | 13 | 19 | 1.055 | 0.108 | 0.069 | **0.003** |

### Behavioral Observations

The methods for gathering behavioral data from these aviaries have been described previously ^43,44^. In brief, expert observers identified birds using distinct colored leg bands, and recorded the actor, receiver, and the behavior type for a variety of vocal and non-vocal behaviors, which were automatically stored with timestamps using speech-to-text software. Observers performed “scan-sampling”, generally attempting to record behavior from as many individuals as possible and from all areas of the aviary rather than focusing on one individual, making this dataset particularly well suited for studying group trends, although it does not capture the full range of behavioral events, nor 100% of song events. Observers could capture roughly one behavior every 1-2 seconds. In some aviaries, as part of other experiments, males or females were added or removed during the breeding season. In these aviaries we only analyzed cohesion up to the day before switches occurred.

In all aviaries, eggs were counted and removed each day. In two aviaries (marked with † in Table 1), individuals were switched prior to the laying of eggs, thus these were excluded from analyses using eggs (e.g., Figure 2–3), although these were used in demonstrating cohesion. In later years, we used CCTV cameras and RFID tags to assign eggs to individual females, however in early experiments, parentage could only be assigned using genotyping (used in the aviaries in Figure 4). For this reason, four of the aviaries (marked with * in Table 1) used to demonstrate the effect of cohesion on overall egg output (Figure 2) were excluded when assessing individual reproductive output (Figure 3).

### Defining Metrics

We tested for the presence of cohesive song behavior using the Two-sample Kolmogorov-Smirnov test for skew performed on the distribution of songs to females within bins across the season, compared to the random distribution. The Kolmogorov-Smirnov test is used to test whether two probability distributions, in this case the observed data versus the random expectation, differ. We generated the distribution of percentage of song to females by binning the behavioral observations into 60s bins. Within each bin, we calculated the proportion of songs (across all males) directed to females. If each male sang one and only one song in every bin, we would expect the random distribution to follow a binomial distribution. In practice, some males sang many songs, while others sang none. To generate a relevant random distribution, we shuffled male behavior, maintaining individual male bins but changing their order. Thus, each male’s average behavior was the same, as was the within-bin behavior, but the order of bins was randomized, and each male’s behavior was independent of each other male. This procedure was repeated 10000 times for each aviary to generate a random distribution, which was tested against the sampled distribution. To ensure cohesion was an effect of coordinated singing strategy per se, and not just an increased probability of simultaneous singing (of any type), we excluded all bins with fewer than 2 males from both the observed and shuffled data. This also guaranteed that every datapoint represented a measure of group cohesion, rather than just individual behavior.

While this previous approach was effective for testing for cohesion within individual aviaries, it was not obvious how to compare the results across aviaries. For this reason, we quantified <u>cohesion</u> as the mean Spearman’s rank correlation coefficient of the percentage of song directed to females. This metric was selected because, unlike many measures of correlated activities (e.g., cross correlation of song events), the correlation of percentage of song to females is not influenced by the overall amount of song. This was particularly important in our dataset, in which song rates vary dramatically between individual birds and between groups, and tend to correlate with dominance, pair bonding, and reproduction.

In testing for cohesion we used 60s bins, to balance the number and size of bins. There was no dramatic effect of bin size, but for the main effect of cohesion and egg score (Figure 2), we calculated cohesion for all bin sizes between 45-90s and computed an average value for each aviary, in order to ensure it was not subject to a bin-size effect. For the juvenile experiment (Figure 4), we used 90s bins to increase the number of songs captured in each bin. For the bar and network plots in Figure 1a-b, behavioral data was divided into 100s bins (in order to maximize the number of songs captured in a single bin for visualization and provide intuitive time ranges).

Within each bin, for each male, we calculated the proportion of songs where a female was marked as the receiver. If the male sang 0 songs, this value was undefined. After calculating the proportion of song directed to females for every male in every time bin, we calculated the pairwise Spearman’s rank coefficient across all overlapping time bins for each male-male pair (thus in rare instances where two males never sang in the same window, their pairwise correlation score was undefined). Cohesion of each aviary (see Figure 1, Figure 2) was defined as the mean correlation across all male-male pairs.

Reproductive score was calculated for each aviary in comparison to the mean expected number of eggs for a given number of females over a given time period. A reproductive score of 2.0 represents twice as many eggs as the expectation, given the number of females and the time period sampled. Because egg production varies across the season, and the date ranges included for each aviary varied (Table 1), we could not simply count eggs per female. Instead, we calculated the mean number of eggs per female produced on any given day in the season, averaging across all aviaries. We smoothed this using a Savitzky-Golay filter (window=11,order=1), with the assumption that the true distribution of eggs is quite smooth over the breeding season. Then, for each aviary we calculated the sum of expected eggs per female across all days included, and then multiplied by the number of females to calculate the expected number of eggs. We divided the actual eggs by this value to get reproductive score. For individual male fecundity, we assigned all eggs produced by his pair bonded female(s), with each female’s pair bond being defined as the male from whom she received greater than 60% of songs.

When visualizing cohesion, it was clear that an individual male’s cohesion (any given row in Figure 1c), could be heavily influenced by a single outlier individual. It did not seem appropriate for a male’s cohesion score to be so influenced by a single strong or weak correlation, thus, to better compare across individuals, individual <u>degree cohesion</u> was defined for each male as the number of correlations greater than .1. We chose this metric to emphasize the number of stronger connections, while also preventing individual outliers from disrupting the measure dramatically. <u>Discord</u> for an individual male was similarly defined as the number of strong negative connections (r < −0.1), while discord for the aviary was the proportion of negative connections, r < −0.1.

### Analyses

All analyses were conducted in Python 3.5 using standard libraries. All statistics were calculated using the *stats* module from scipy in python, with the exception of mixed effect models, which were performed using the *statsmodels* python module. We used custom python code for bootstrapping confidence intervals and for averaging across window size (Figure 2). Network figures and analyses were produced using *NetworkX*, with post-processing to alter the shape used to depict juveniles and position of the females (placing them adjacent to their respective pair bonds) using Affinity Designer. No other post-alterations were made to networks. All figures were created using *Matplotlib*, with minor formatting edits using Affinity Designer. All source data, along with the code for all analyses as a Jupyter notebook, are available on GitHub (github.com/aperkes/aviaryanalysis)

---

**Supplemental Figure 1.**
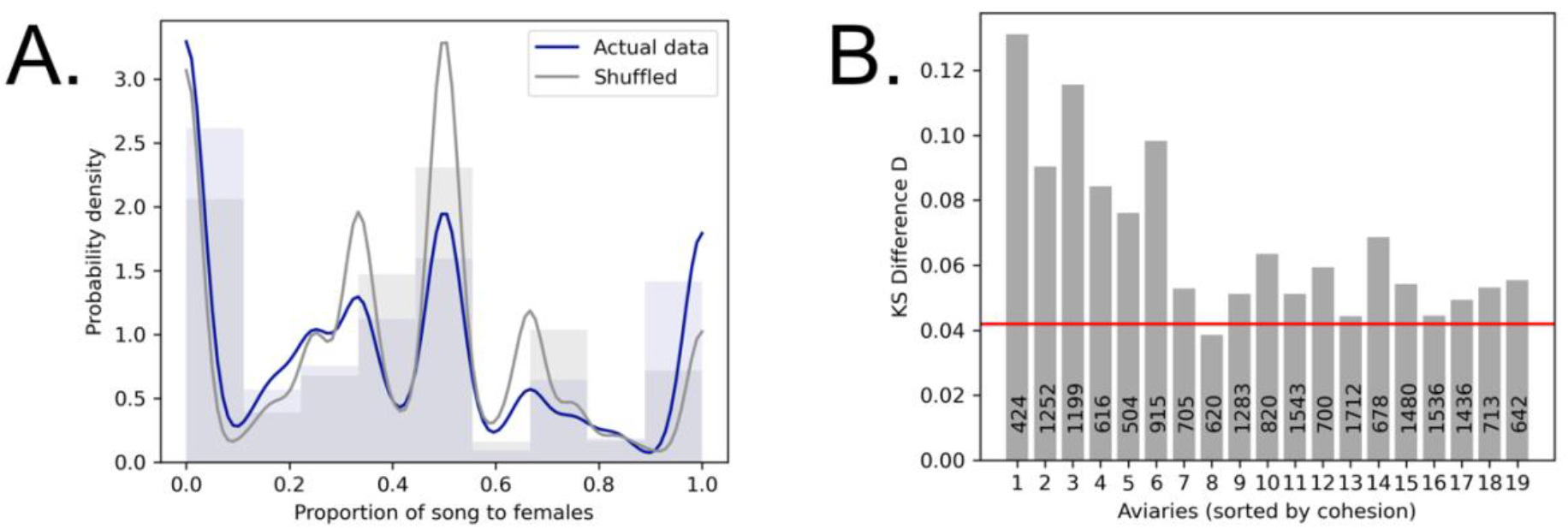
Aviaries display statistically significant cohesion. **A. Distribution of song to females for observed and shuffled data.** This density plot shows the real versus shuffled distribution for the proportion of songs sung to females. The actual distribution was produced by dividing the breeding season into 60s bins and summing the songs across all males. The shuffled distribution was produced similarly, but the bins for each male were shuffled randomly (and this procedure was repeated 10,000 times). Here, the actual data is significantly different from chance (KS Test, D=0.06,p<0.0001). Note the increased instances of 100% female song and 0 % female song (representing 100% male song) in the real data compared to what would be expected by chance, reflecting cohesive behavior. **B. Cohesion across all aviaries.** This plot shows the results of the Kolmogorov-Smirnov test, calculated as described in A. for each aviary. All but one aviaries are significantly more cohesive than their random distributions. The red line represents the significance at n=1050 and is placed so as to divide the 18 significant aviaries from the one non-significant aviary. Actual sample sizes vary, and are written on each bar, representing the number of bins (from the real data) in which at least 2 males sang song of any type (see methods for full details). Aviaries are sorted as in Figure 1e.

