## Supplementary figures and images for "Cohesion in male singing behavior predicts group reproductive output in a social songbird"

### Supplemental Figure 1

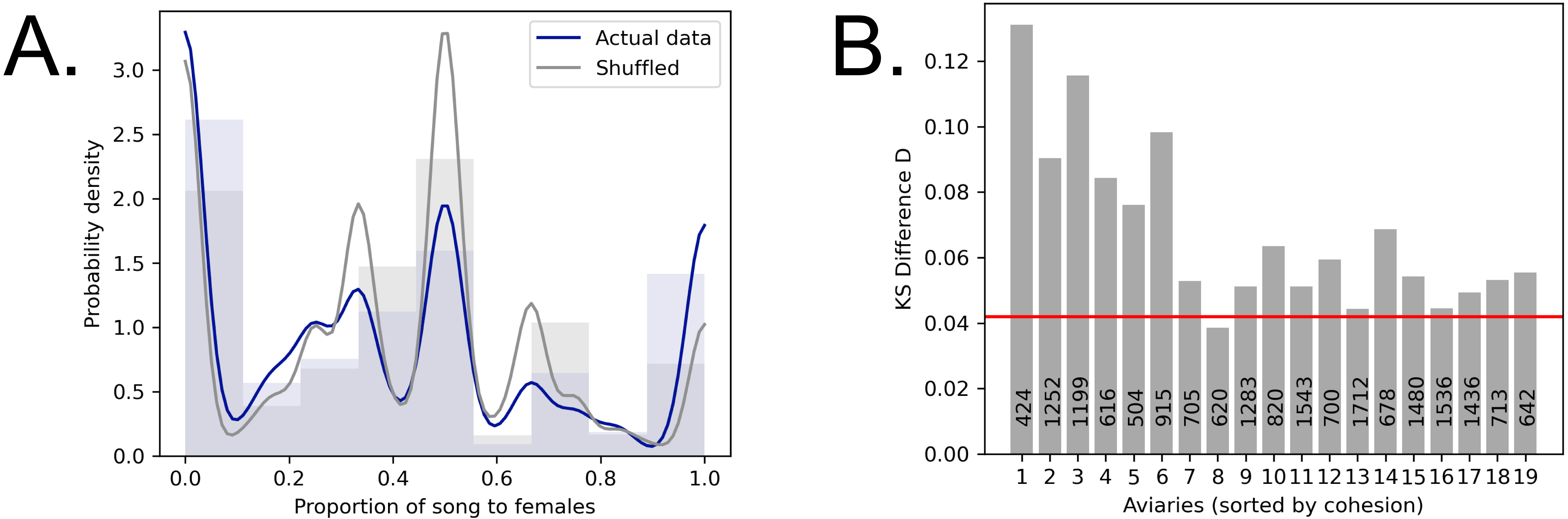
